# Overlapping Cortical Substrate of Biomechanical Control and Subjective Agency

**DOI:** 10.1101/2024.07.24.604976

**Authors:** John P. Veillette, Alfred F. Chao, Romain Nith, Pedro Lopes, Howard C. Nusbaum

## Abstract

Every movement requires the nervous system to solve a complex biomechanical control problem, but this process is mostly veiled from one’s conscious awareness. Simultaneously, we also have conscious experience of controlling our movements—our sense of agency (SoA). Whether SoA corresponds to those neural representations that implement actual neuromuscular control is an open question with ethical, medical, and legal implications. If SoA is the conscious experience of control, this predicts that SoA can be decoded from the same brain structures that implement the so-called “inverse dynamics” computations for planning movement. We correlated human (male and female) fMRI measurements during hand movements with the internal representations of a deep neural network (DNN) performing the same hand control task in a biomechanical simulation– revealing detailed cortical encodings of sensorimotor states, idiosyncratic to each subject. We then manipulated SoA by usurping control of participants’ muscles via electrical stimulation, and found that the same voxels which were best explained by modeled inverse dynamics representations— which, strikingly, were located in canonically visual areas—also predicted SoA. Importantly, model-brain correspondences and robust SoA decoding could both be achieved within single subjects, enabling relationships between motor representations and awareness to be studied at the level of the individual.

**Significance Statement:** The inherent complexity of biomechanical control problems is belied by the seeming simplicity of directing movements in our subjective experience. This aspect of our experience suggests we have limited conscious access to the neural and mental representations involved in controlling the body – but of which of the many possible representations are we, in fact, aware? Understanding which motor control representations percolate into awareness has taken on increasing importance as emerging neural interface technologies push the boundaries of human autonomy. In our study, we leverage machine learning models that have learned to control simulated bodies to localize biomechanical control representations in the brain. Then, we show that these brain regions predict perceived agency over the musculature during functional electrical stimulation.

## Introduction

Even simple movements require the nervous system to solve a complex control problem. The human hand alone has more than 20 degrees of freedom (i.e., directions joints can move) and is controlled by over 30 muscles. Complicating the problem, each joint can be moved by multiple muscles, each of which move multiple joints (Bullock et al., 2012). Theoretical accounts posit our brains contain a vast, interacting set of internal representations of the body’s positional states and kinetics, but little of this machinery is accessible to awareness (Blakemore et al., 2002). The act of moving *feels* straightforward and is accompanied by a sense of agency (SoA)—a feeling of “I did that.” Does this conscious experience of directing movement reflect the actual machinery of control, and if so, what types of motor representation influence awareness?

Most extant research focuses on *prediction errors* between expected and actual sensory outcomes as determinants of SoA (Frith, 1987; Synofzik et al., 2008). However, not all errors diminish SoA—some mismatches can even overwrite our original intention in memory (Lind et al., 2014). Since errors only account for *negative* experiences of agency, not why we feel SoA in the first place, such “comparator” models fall short of predicting which mismatches are consciously detected (Frith, 2012).

The need for a predictive account of SoA is becoming urgent. Advanced neuroprostheses leverage the technology behind today’s large language models to “autocomplete” instructions decoded from the brain (Metzger et al., 2023; Tang et al., 2023). These interfaces may pose a threat to the autonomy of those they aim to help if deviations from user intentions are not accessible to awareness. Some countries have already passed legislation to this effect (Fernández and Fernández, 2022). However, without a foundational understanding of which motor representations affect consciousness, it is unclear how to design neural interfaces around these constraints.

Normative theories of motor control posit two sorts of internal models: *forward dynamics models* to predict future states of the body from current neuromuscular output, and *inverse dynamics models* or *control policies* to generate neuromuscular output that achieves a desired state (Wolpert and Ghahramani, 2000). By some accounts, paired forward and inverse models are specialized for specific *sensorimotor contexts* (e.g. moving with rested vs. fatigued muscles), and forward model prediction errors are used not just to update the models for the current context, but also to identify when contexts switch (Wolpert and Kawato, 1998). The recent COIN model (Heald et al., 2021), which posits neuromuscular output is averaged over all control policies (weighted by the probability of being in each context) but subjects only have awareness of one context at a time, quantitatively predicts the contribution of explicit (i.e. conscious) and implicit components of adaption following rotation of subjects’ visuomotor mapping. Following this logic, SoA may persist following a prediction error if the error can be explained by another sensorimotor context for which one already has a control model, but SoA is lost when one lacks an inverse dynamics model for the current best-guess context.

This hypothesis makes a testable prediction that SoA can be *decoded* from the same cortical areas where representations of an inverse dynamics model are putatively *encoded* (in the context for which that model would ordinarily apply). In the present study, participants performed hand gestures during functional MRI. To localize inverse dynamics representations, we predicted participants’ voxelwise brain activity during the task from activations of a deep neural network (DNN) that performed the same task with a simulated biomechanical hand (Caggiano et al., 2022). Then, in another session, we usurped control of subjects’ muscles using functional electrical stimulation; by slightly preempting subjects’ endogenous movements, we elicit an erroneous SoA over roughly half of involuntary movements as validated in our previous work (Veillette et al., 2023). We test whether we can decode participants’ SoA over individual muscle movements from those same voxels that were best predicted by inverse dynamics representations days prior.

## Methods

### 1. Participants and Ethics Statement

Four participants (3 female, 1 male), between the ages of 24 and 28 years, participated in the study. All subjects gave their written, informed consent before participating, and all procedures were approved by the Social and Behavioral Sciences Institutional Review Board at the University of Chicago (IRB23-1323).

### 2. Experimental Design

#### 2.1. Session 1: Motor Task

Upon arriving at the MRI facility, subjects put on an MRI-compatible Data Glove (5DT, Inc.) which was used to measure their joint angles throughout the subsequent recording. The Data Glove only outputs raw sensor values for each joint instead of joint angles directly, though these uncalibrated sensor values linearly vary with to the true joint angles. To calibrate the glove, we recorded participants moving their gloved hand outside of the scanner with a Leap Motion Controller (Ultraleap), from which we estimated ground truth joint angles using the *RoSeMotion* software (Fonk et al., 2021). We used this data to estimate the linear mapping from the glove sensors to true joint angles (including the distal interphalangeal joints for which the glove does not have direct sensor coverage) on a per-subject basis.

Subjects were instructed to replicate hand positions that were presented to them as pictures of a hand (Avotec Silent Vision SV-6011 LCD projection system), which switched every five seconds while in the MRI scanner. As stimuli, we used the same eight isometric and isotonic hand configurations used in the *Ninapro* database (Jarque-Bou et al., 2020). Over 120 trials, we exhausted each of the 56 possible transitions between the gestures at least twice during each of the ten, 10-minute runs. During this, we recorded their joint angles and velocities using an MRI-compatible motion tracking glove (60 Hz sampling rate). While target positions were presented visually, participants could not see their own hands outside of the scanner. The unit of analysis was not these 5-second trials, but prediction of the blood oxygen level dependent (BOLD) measurement at each voxel in each fMRI measurement (every 2 seconds). Overall, this session lasted 2.5 hours. Data from this task was used to fit voxelwise encoding models designed to capture the inverse dynamics computations thought to be necessary for musculoskeletal control (see Fig. 1).

**Figure 1:**
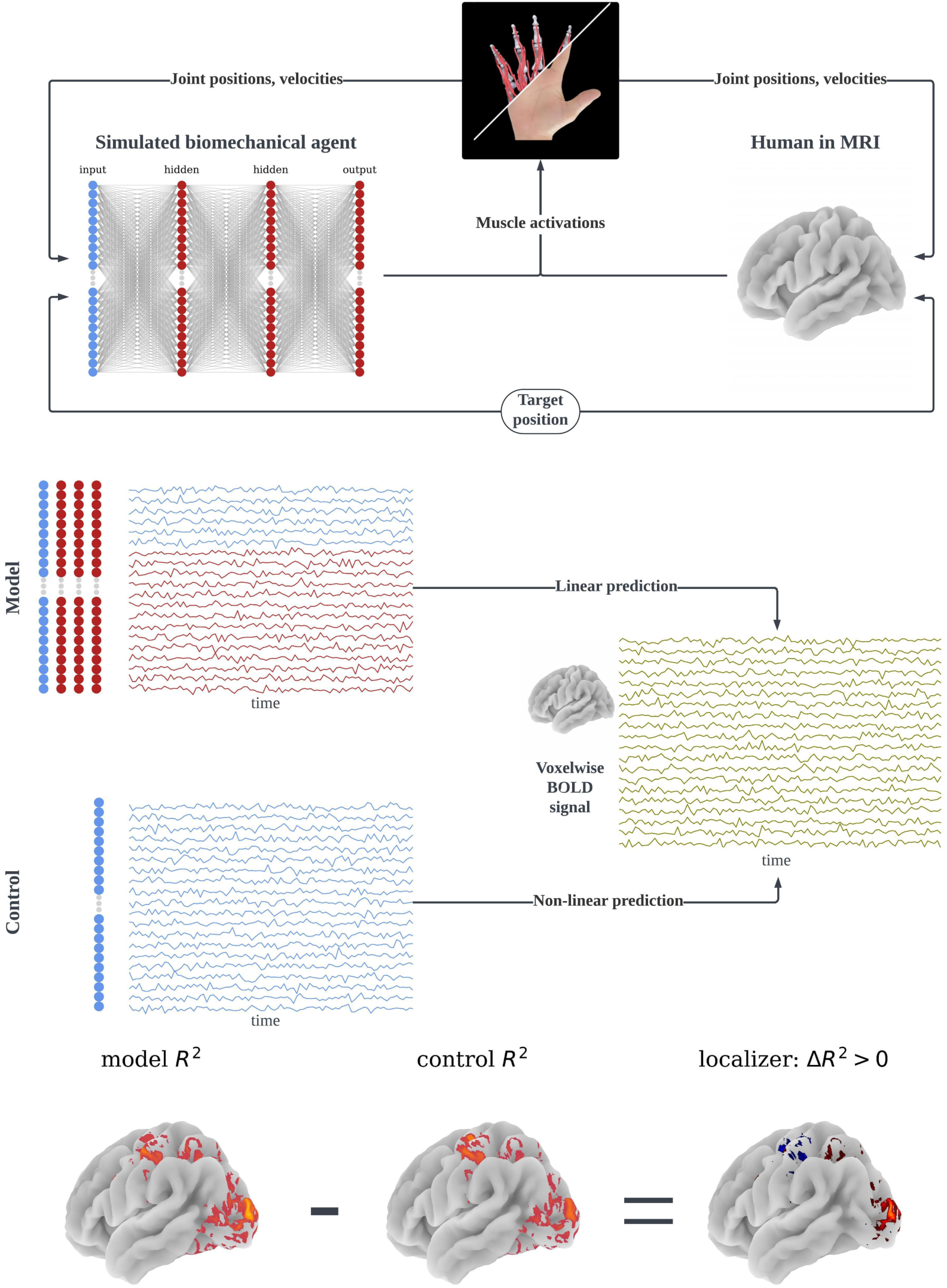
Approach to localizing inverse dynamics model representations in cortex. Subjects moved their hands in specified “target positions” while we recorded their hand movements in an fMRI scanner. We used the activations of a deep neural network (DNN) which performs the same hand pose task with a simulated biomechanical hand—i.e., approximates an “inverse dynamics model” or controller for the hand—as features to predict voxelwise brain activity over time. We compare the out-of-sample predictive accuracy of the inverse dynamics DNN model to a purely data-driven voxelwise encoding model, and the voxels that are better predicted by our biomechanics-informed model are interpreted as reflecting the representations of an inverse dynamics model.

#### 2.2. Session 2: Agency Task

In a second session, participants returned to complete ten 10-minutes runs of a cue-response reaction time task, pressing a button with their ring finger as quickly as possible following a visual prompt. After the first (baseline) block, we applied functional electrical stimulation (FES) to the *flexor digitorum profundus* muscle to produce a muscle movement which would cause an involuntarily press of the button around the time participants would respond on their own. After each trial, they were asked to discriminate whether they or the muscle stimulator caused the button press. Using a Bayesian adaptive procedure developed in our prior work (Veillette et al., 2023), we continuously adjusted the stimulation latency until subjects responded that they caused the button press ∼50% of the time (see Figure 2a)—which, for most subjects, is robustly before they could have possibly begun to move (Kasahara et al., 2019; Tajima et al., 2022; Veillette et al., 2023). Trial-by-trial agency judgements in this task can be decoded from electroencephalogram (EEG) within the first 100 ms following the onset of muscle stimulation, indicating that variation in self-reports usually reflects sensorimotor processes rather than post hoc guessing (Veillette et al., 2023).

**Figure 2:**
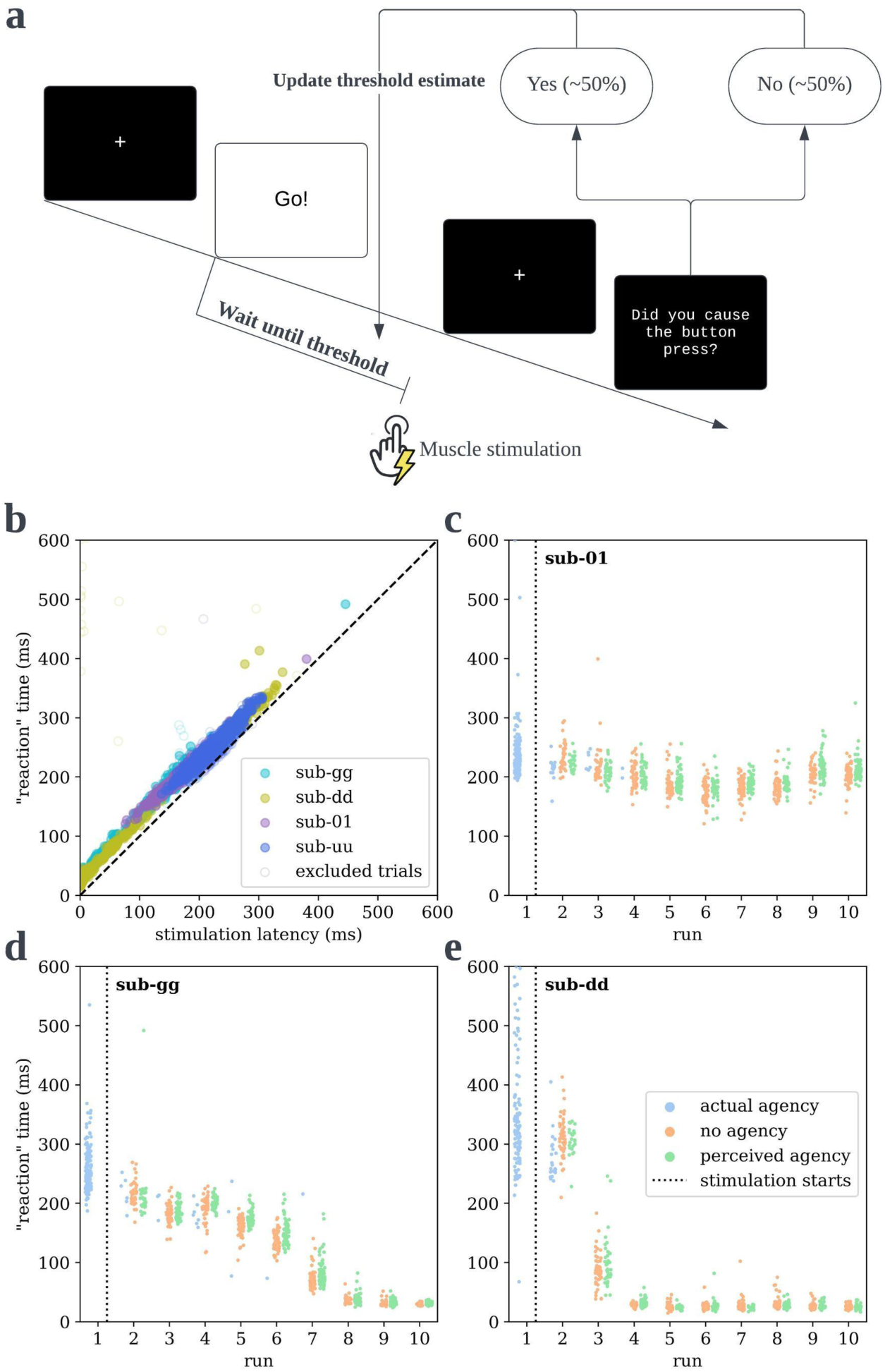
Manipulating sense of agency (SoA) by usurping control of subjects’ muscles with electrical stimulation. (a) Subjects complete a cue-response reaction time task, but we used muscle stimulation to preempt their self-produced movements on most trials. Subjects were asked to discriminate, after each trial, whether they or the muscle stimulation caused their finger to press the button. Their response was used to update a threshold estimate on which the timing of stimulation was based, such that we could guarantee they believe they caused roughly 50% of movements. (b) As expected from our prior work, this 50% threshold is substantially earlier than subjects move autonomously, as illustrated by the fact that button presses (“response” times) are essentially a linear function of stimulation latency—seen here for all stimulation trials across all subjects. (c-e) This adaptive procedure tracks non-stationarity in subjects’ threshold, such that subjects rarely move faster than the muscle stimulator (blue) after the first run with stimulation. Consequently, we obtain distributions of “no agency” and “perceived” but false agency trials with highly overlapping stimulation latencies, so subjects’ experienced SoA can be dissociated from the task manipulation.

Upon subjects’ arrival, we applied two functional electrical stimulation (FES) electrodes to their forearm over the *flexor digitorum profundus* muscle, which were connected to a RehaStim 1 constant current stimulator (HASOMED) through a waveguide. Before the experiment, we ran a calibration procedure in which we raised the intensity of the FES stimulation until it could cause subjects’ ring finger to press a button (Celeritas optical response pad) ten times in a row to ensure we could adequately move their muscles with FES. Each instance of stimulation consisted of 3 consecutive, 400 microsecond (200 pos, 200 neg) biphasic pulses.

The task procedure was the same as used in our prior research (Veillette et al., 2023), but with more trials spread across ten blocks. The first 10-minute run was a typical cue-response reaction time task, in which subjects were asked to press the response pad with their ring finger as quickly as possible after they see a cue to move. In the remaining 9 runs, subjects were instructed to still attempt to complete the reaction time task on their own, but if they were not fast enough, the muscle stimulator would move their finger to press the button for them. If subjects succeeded in moving before FES, it would trigger stimulation immediately so the muscle movement and FES were always temporally confusable. After each trial, subjects were asked to discriminate whether they or the muscle stimulator pressed the button. (If they were unsure, as it can be surprisingly difficult to discern, they were told to provide their best guess.)

We used subjects’ response times from the first run to set a prior on the parameters of a logistic function describing the probability that they would report causing the button press, based on the observation from previous work that FES-caused movements can occur up to 40-80 ms prior to self-caused movements before subjects notice the have been preempted more than half the time (Kasahara et al., 2019). After each trial, we used their agency judgments to update this model, and we draw the next stimulation latency from the posterior distribution of threshold at which they would report agency or non-agency with equal probability. To account for nonstationarity between runs, the uncertainty in the posterior is reset at the beginning of each run, though the posterior mean is retained.

As we know from our prior research (Veillette et al., 2023), this Bayesian adaptive procedure produces distributions of agentic and non-agentic trials with very similar distributions of stimulation latencies—so that participants’ subjective experience of agency can be dissociated from the stimulation parameters. In practice, this 50%-50% threshold is sufficiently early that subjects are rarely able to respond before the stimulator (see Figure 2c-e), which can be verified by checking that measured “response” times are a linear function of FES latency (see Figure 2b), and those trials that do not follow the line can be removed following the criteria in our prior work (Veillette et al., 2023). In the present study, this yielded 877, 853, 926, and 728 FES-caused trials across the nine stimulation blocks for subjects 01, *dd*, *gg*, and *uu*, respectively, that we used for our decoding analysis. Only FES-caused trials were used for decoding, as other, unobserved aspects of the muscle movement may differ between FES-caused and self-caused movements.

Additionally, we can rule out the possibility that self-reported agency judgments at this threshold latency are just random, as we have found that single trial judgements can be decoded (cross-validated across subjects) from EEG within the first 100 ms after the onset of muscle stimulation and remain decodable for at least another 400 ms, indicating that the agency judgments in this task usually have an origin in sensorimotor processing (Veillette et al., 2023). Of course, even if the average subject responds nonrandomly, some subjects might still have very high guess rates and be effectively random. This outlier case can be identified if subjects’ responses are insensitive to FES latency (see analysis).

An unanalyzed 10-minute eyes-open resting state scan was collected during this session after the fourth run, which was there merely to serve as a brief break for the subject but is available with our open dataset. The experiment session lasted 2.5 hours in total.

### 3. Statistical Analysis

#### 3.1. Voxelwise Encoding Models

To build our biomechanics-informed encoding model, we used a deep neural network (DNN) that was trained to generate muscle activity that would shape a simulated hand into specified hand gestures, like our human participants, in a biomechanical simulation. Specifically, we used the pretrained model released with *MyoSuite’*s hand pose simulation environment (*MyoHandPoseRandom-v0*); this model maps a current task state (i.e., joint angles, velocities, and distance from target hand position) to a set of muscle activations that aim to achieve the target position (Caggiano et al., 2022). This deep neural network (DNN) approximates an inverse dynamics model, as it takes a current state as input and outputs a set of muscle activations that will move its biomechanically realistic hand closer to the target position. Though it does not reach human-level performance—and importantly, it never saw human data during training—the model achieves a joint angle error (summed across joints and simulation time) of 1.94 radians after 1,000 training iterations (compared to 3.29 rad after the first training iteration) using the natural policy gradient method (Kakade, 2001).

To translate this approximation into a neural encoding model, we input each measurement of our human participants’ task state—position and velocity via the motion tracking glove, as well as the target position—at each time point into the DNN and extracted all activations from all artificial neurons; these activations were used as features to linearly predict the subjects’ continuous BOLD activity using a separate ridge regression for each voxel (see Figure 1).

These voxelwise encoding models were fit for each participant using the *Himalaya* package (Dupré la Tour et al., 2022). Features from the DNN were filtered to the same rate as the MRI data, and then duplicated with four temporal delays (2, 4, 6, and 8 seconds) to account for the lag between neural activity and the hemodynamic response. A separate linear ridge regression was fit for each voxel, resulting in 652 weights (4 times 163 DNN units) for each voxel, were averaged across delays to produce a 163-dimensional vector describing each voxel’s response properties. The regularization parameter for each ridge regression was chosen by grid search to maximize the leave-one-run-out cross-validated 𝑅^2^ within the training set.

To account for the possibility that the task features, which include the participant’s actual movements, alone drove prediction rather than inverse dynamics features, a separate control encoding model was fit similarly but only had access to the input features of the neural network as predictors. Instead of linear ridge regression, this encoding model used kernel ridge regression such that it could also learn nonlinear mappings between task features and voxel activity. This ensures that, when our DNN-based encoding model outperforms the control, it is because the inverse dynamics features are *particularly* good features for predicting brain activity (i.e., better than a data-driven nonlinear mapping)—not because any arbitrary nonlinear transformation of the input space improves performance.

Encoding models were fit to the first seven MRI runs, and then tested (cross-validated) on three hold-out runs which used a separate MR field map measurement such that fMRI preprocessing for these runs was totally independent. For group-level analyses, these cross-validation scores were averaged across participants. Models’ out-of-sample 𝑅^2^ were compared to chance using a block permutation test (5,000 permutations) in which continuous blocks of 10 TRs are kept together on each permutation so that the autocorrelation structure of the data is preserved in the null distribution (Huth et al., 2016; LeBel et al., 2023). In visualizations of encoding model performance (before subtracting the control model’s performance) and in the “visuomotor mask” used for decoding, we apply a false-discovery rate (FDR) correction and keep voxels where FDR < 0.05, as in other voxelwise encoding studies (Huth et al., 2016; LeBel et al., 2023; Tang et al., 2023). In-text, we report the lowest familywise error rate corrected *p*-value across the brain as a “global” *p*-value for each subject, corrected for multiple comparisons using an 𝑅^2^-max procedure (Nichols and Holmes, 2002). When comparing the DNN-based and control models, we use a paired block permutation test, in which blocks of model predictions are shuffled between models rather than in time, and we control the familywise error rate using threshold-free cluster enhancement (Smith and Nichols, 2009).

To visualize DNN-based encoding models, we subdivided the inputs of the DNN into three features spaces, and we computed Shapley values using the *DeepLIFT* method with the *shap* package (Lundberg and Lee, 2017; Shrikumar et al., 2017). Shapley values describe how much each feature or feature space contributes to the predictions of a model or, roughly, how much the model’s predictions (in our case, of a voxel) would change if features were not included. For visualization (depicted in Figure 3), we show each feature space’s test-set Shapley values divided by the sum of the Shapley values for all feature spaces, so each voxel’s color represents the relative (i.e. proportional) contribution of each feature space for explaining that voxel.

**Figure 3:**
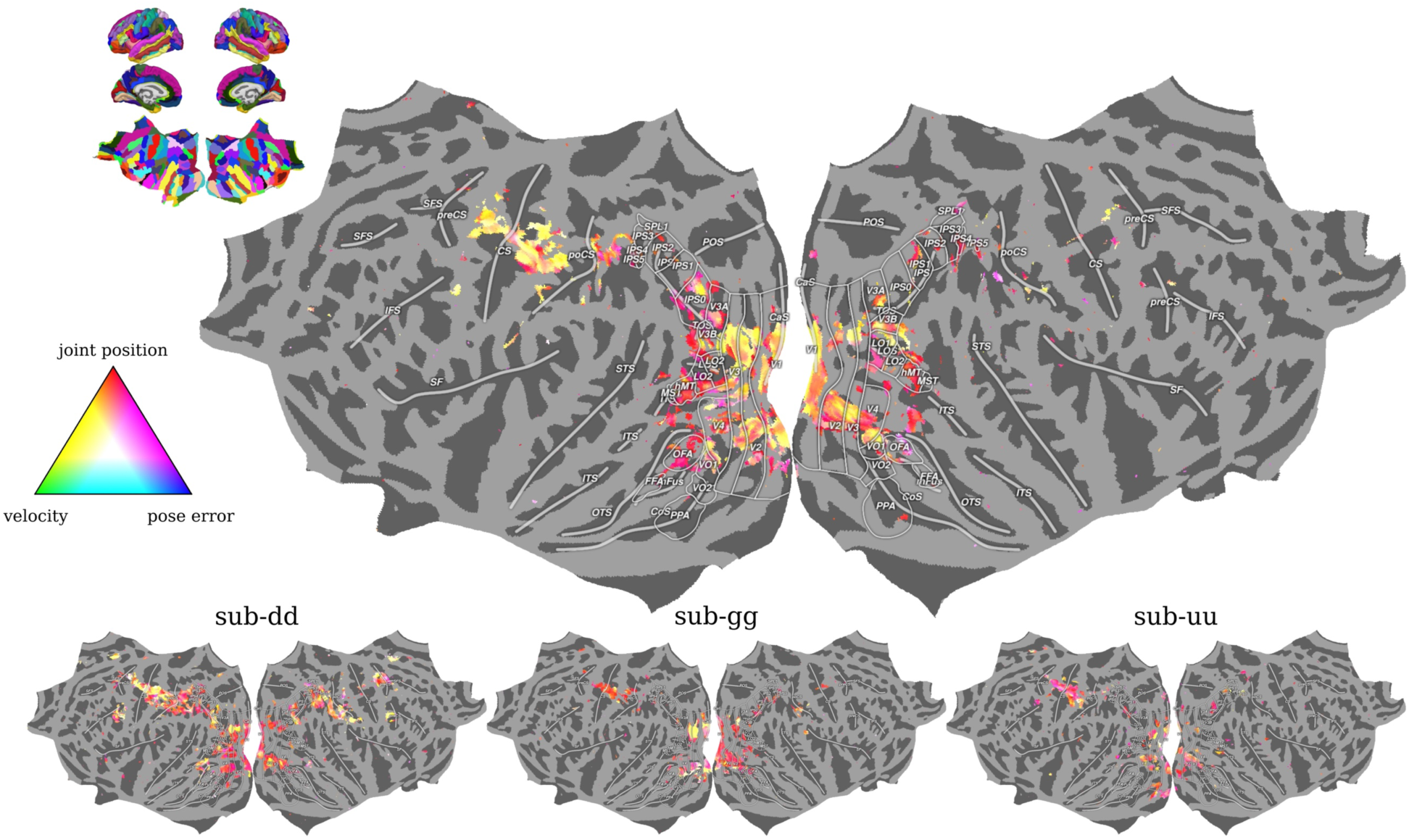
Voxelwise response properties while moving the hand. Relative importance of sensorimotor task state features during the motor control task were estimated using Shapley values. Each voxel that is significantly predicted by the DNN-based encoding model (FDR < 0.05) is assigned an RGB color based on these feature importances. For example, a voxel colored red is predicted by DNN units that respond exclusively to hand position, but a voxel colored pink behaves like DNN units that respond to combinations of hand position and deviance from target position—i.e., likely encode the target position.

#### 3.2. Decoding Models

For decoding, voxel activity for each trial was estimated by computing the beta series for the FES stimulation events using the “least squares all” method (Rissman et al., 2004). We used logistic regression with lasso (L1) regularization to predict participants’ single trial agency judgments (from the agency task) from the full-rank set of principle components of the voxel activity—called logistic lasso-PCR. Lasso-PCR is commonly used for whole-brain decoding models, as the principle components transformation usefully deals with the high spatial autocorrelation of fMRI measurements, and thus the method scales quite well with the number of voxels included as predictors—but the PCA transformation can be inverted to easily project model weights back into interpretable voxel space (Wager et al., 2011).

We fit decoding models using three nested feature sets: (1) our “theory mask,” consisting of those voxels the inverse dynamics DNN model predicted better than the control, (2) a “visuomotor model” consisting of all the voxels predicted above-chance by the control model, and (3) all the voxels in cortex. Models performance was quantified as the leave-one-run-out cross-validated area under the receiver operator characteristic curve (AUROC), which is interpreted similarly to accuracy (0.5 is chance, 1 is perfect, 0 is worst) but is not dependent on a threshold criterion and thus invariant to “how much” agency a subject must feel before claiming they caused a movement—and is usefully robust to class imbalances. Group-level analyses use the mean AUROC across all participants; we report “theory mask” decoding results using both the mask derived group the group-level analysis as well as individual-specific masks if available (i.e. in which the inverse-dynamics-based encoding model significantly outperformed the control model for that individual participant).

Models are compared to chance (i.e., at an AUROC of 0.5) using a one-tailed permutation test. Nested models are compared to each other using a one-tailed paired permutation test. (A one-tailed test is used when comparing models with nested feature sets, since it is assumed that all the information captured by the smaller set of models is also present in the larger set, and if the larger set underperforms the only explanation is theoretically uninteresting overfitting with the larger feature set.)

Stimulation latency is the means by which we manipulate subjective agency. Thus to account for the possibility that we might just be decoding stimulation latency, we ran a logistic regression (with nested random effects for each participant and BOLD run) predicting agency judgments from (a) the stimulation latency and (b) the out-of-sample prediction of the “theory mask” model for each trial. To ensure robustness against violations of normality assumptions (e.g., as a biproduct of cross-validation), we fit this logistic regression using generalized estimating equations (GEE), which is more robust to violations of distributional assumptions than is maximum likelihood estimation (Liang and Zeger, 1986). Since this model statistically controls for the stimulation latency, we can interpret a nonzero coefficient assigned to the brain-based prediction as evidence that the decoder captures variation in sense of agency that is *not* explained by the stimulation latency. Additionally, the coefficient for the stimulation latency quantifies the subjects’ sensitivity to the timing of their muscle movement around their threshold, or the stability of their threshold over time.

#### 3.3. Within-Subject Analyses

Rather than testing a large number of participants for a short amount of time as in typical fMRI studies, we focused on collecting sufficient amounts of data to establish robust model-brain correspondences in each of four participants (i.e., 3.5 hours of BOLD data, collected over 5 hours). By focusing on the individual rather than the group as the unit of analysis, this “dense sampling” approach can be more sensitive to neural patterns that are robust within individuals but idiosyncratic across them (Poldrack, 2017; Naselaris et al., 2021), which is critical for modeling neural encodings of high-dimensional feature spaces (Cross et al., 2021; Tang et al., 2023). This has led some researchers to advocate treating each participant as their own *n* = 1 experiment and estimating the population prevalence, rather than the group mean, of experimental effects (Allefeld et al., 2016; Hirose, 2021; Ince et al., 2021, 2022; Veillette and Nusbaum, 2025).

To this end, we also performed our main analyses at the individual level, treating each subject as an *n* = 1 experiment. When reporting single-subject results, the first subject is labelled “sub-01” and other subjects are given arbitrary letter codes, to denote that they are replications of the first *n* = 1 experiment. For each analysis, we report a *p*-value for each subject (with their subject ID as a subscript), and we combine *p*-values across subjects meta-analytically using Fisher’s method—this combined *p*-value (denoted 𝑝_all_) corresponds to the null hypothesis that there is no effect in any subject, accounting for multiple comparisons. A significant result using Fisher’s method indicates some people in the population show an, but it does not imply the effect is representative—a claim which would require explicit prevalence inference.

#### 3.4. Prevalence Inference and Sample Size Considerations

Despite the increasing popularity of small-*n*, densely-sampled designs such as our own, there is no consensus concerning how many participants are “enough” for such a study to make general claims. However, several criteria have been proposed by various research groups, which all relate back to the question of how many individual participants must be seen to show an effect before concluding that effect is present in a substantial portion of the population.

Replying to an open review of their influential recent paper (Gordon et al., 2023), Dosenbach and Gordon suggested that “precision neuroimaging” studies should adopt the conventions of the nonhuman primate literature, in which densely sampling from a small number of subjects has long been the norm (Dosenbach and Gordon, 2023). It is convention in that literature to report a finding only when at least 2-out-of-3 or 3-out-of-4 primates in the study show an effect individually; simulations suggest that following either such a rule-of-thumb is exceedingly unlikely to yield an effect that is not present in a non-negligible portion (i.e. >20%) of the population (Laurens, 2022). This is the rule we initially used to select our sample size, using the 3-out-of-4 rule such that, should only 3 subjects show significance in the localizer session, we could still us the 2-out-of-3 rule for analysis in the second session.

A less *ad hoc* framework for sample size determination has recently been proposed (Schwarzkopf and Huang, 2024). Noting that the probability of a within-participant hypothesis test (i.e. *n* = 1 experiment) with Type I error rate 𝛼 and Type II error rate 𝛽 yielding a statistically significant result is simply 𝛼 if the null is true for the whole population (all-𝐻_0_) and is 1 − 𝛽 if instead the alternative is always true (all-𝐻_1_), Schwarzkopf and Huang (2024) propose testing between the all-𝐻_0_ and all-𝐻_1_models using a sequential probability ratio test (SPRT) (Wald and Wolfowitz, 1948). In Wald’s SPRT, a researcher recomputes the likelihood ratio of two models, Binomial(𝑛, 1 − 𝛽)/Binomial(𝑛, 𝛼) in our case, after every observation (i.e. participant) and stops data collection once this ratio crosses either an upper threshold (in which case we would conclude in favor of all-𝐻_1_) or lower evidence threshold (concluding all-𝐻_0_). Thus, sample size is determined adaptively, and according to the bounds Wald derived, using the thresholds [0.053, 19.00] contains both the Type I and Type II error rates of this procedure to 5% with optimal sample efficiency (Wald and Wolfowitz, 1948). Since the probability ratio itself depends on the sensitivity 1 − 𝛽 of the within-subject hypothesis test, Schwarzkopf and Huang (2024) recommend simply reporting the range of sensitivities (i.e. power) over which the SPRT would have concluded in favor of the all-𝐻_1_ model as evidence for a claim; this approach is conceptually similar to reporting the smallest effect size one’s sample is well-powered to detect in a fixed-sample design (Lakens, 2022). For our main results (whole-brain significance of encoding model contrast and “theory mask” decoding model), we report the range of sensitivities for which the SPRT would have stopped data collection at or before our sample size to demonstrate we meet this evidence criterion.

We think Schwarzkopf and Huang’s (2024) criterion—deeming a sample size sufficient when it is substantially more likely that the whole population shows an effect than that nobody shows the effect—is reasonable for the aims of this kind of small *n* study, where each individual is so deeply sampled that each subsequent participant is very costly. (Indeed, a single participant in the present study costs as much as five participants in most fMRI studies with hour long sessions.) However, we note two major problems with their approach: (1) Binarizing participant-level data into the accept/reject decisions of a within-participant hypothesis test makes the likelihood ratio in the SPRT highly sensitive to minor deviations in the data when within-participant *p*-values are near the significance threshold, (2) their use of SPRT considers only cases where all participants either do or do not show an effect, *a priori* not considering the possibility that only some participants show an effect, and (3) effect size information present in the data is not used to update the likelihood of the data under 𝐻_1_, making the approach fully reliant on *a priori* assumptions about whether the effect size (i.e. power) falls in the reported range. To overcome these limitations, we recently proposed Bayesian *p*-curve mixture models (Veillette and Nusbaum, 2025), which can calculate a Bayes Factor quantifying the relative evidence for all-𝐻_1_vs. all-𝐻_0_ models of the population (denoted here as 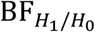) from a series of within-participant *p*-values from repeated *n* = 1 experiments. Unlike the SPRT approach, it can also quantify evidence for a mixture model in which some participants show the effect and others do not vs. for those homogenous population models (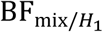 and 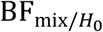). And usefully, the interpretation of a Bayes Factor is invariant to the stopping rule used to determine sample size (Rouder, 2014). The *p*-curve mixture integrates over posterior uncertainty about the within-participant effect size/sensitivity, thus yielding a single evidence ratio per comparison. We report these Bayes Factors for our main analyses, and when a mixture model is most supported given the data, we also use *p-*curve mixtures to estimate the posterior mean and 95% highest density interval (HDI, or credible interval) of the prevalence of 𝐻_1_, which is the proportion of the sampled population expected to show an effect. Bayes Factors and posterior distributions are estimated using the *p2prev* Python package, which we describe and validate in a separate publication (Veillette and Nusbaum, 2025). To facilitate interpretation, we note that by convention Bayes Factors greater than 3 are interpreted as moderate evidence and Bayes Factors above 10 are interpreted as strong evidence (Kass and Raftery, 1995)—though we also note the arbitrariness of these estimates.

Notably, while a group-level effect is often taken to imply that an effect is typical in the population, that is a misinterpretation. Even (actually especially) when the number of participants is large, the group mean may significantly differ from the null even when the prevalence of the effect in the population is very small (Allefeld et al., 2016; Hirose, 2021; Veillette and Nusbaum, 2025). As such, formal prevalence inference based on participant-level results provides complementary information to group-level analyses, so we report both for our main findings.

### 4. fMRI Acquisition and Preprocessing

MRI data were collected with a 3T Philips Achieva at the Magnetic Resonance Imaging Research Center at the University of Chicago. Functional scans were collected using gradient echo EPI with TR = 2.00 s, TE = 0.028 s, flip angle = 77 deg, in-plane acceleration (SENSE) factor = 2, voxel size 3.13×3.13×3.0 mm, matrix size = (64, 62) with 32, FOV = 200 mm. FOV covered cortex in its entirety in all subjects. Pepolar field maps were collected between every 2-4 functional scans to be used for susceptibility distortion correction. High-resolution (1×1×1 mm) anatomical scans were collected during Session 1 on the same 3T scanner with a T1-weighted MP-RAGE sequence.

Data was first preprocessed with fMRIPrep 23.2.0, which was used to perform susceptibility distortion correction, slice time correction, brain extraction, co-registration of functional and anatomical scans, computation of confound (e.g., motion) time series, cortical surface reconstruction, and projection of BOLD data onto the Freesurfer *fsaverage* template surface, in which subsequent analyses were performed (Fischl, 2012; Esteban et al., 2019). CompCor confounds generated by fMRIPrep were regressed out of the raw BOLD time series data (Behzadi et al., 2007) using the *nilearn* package, either prior to fitting voxelwise encoding models for the motor control task or as part of the estimation of beta series (see Methods 3.2) for the agency task.

### 5. Data and Code Availability

Anonymized raw data and preprocessed derivatives, including detailed data quality reports and the fitted encoding/decoding models, have been made publicly available on *OpenNeuro* (https://doi.org/10.18112/openneuro.ds005239.v1.0.1). The experiment code used for data collection for the hand tracking task (https://doi.org/10.5281/zenodo.12610625) and the muscle stimulation task (https://doi.org/10.5281/zenodo.12610710) are archived on *Zenodo,* as well as all analysis code (https://zenodo.org/doi/10.5281/zenodo.12610621).

## Results

### Encoding of sensorimotor states across cortex is widespread but idiosyncratic

Our voxelwise encoding models describe each voxel’s encoding properties as a 163-dimensional vector containing linear model weights for each neuron in the biomechanics-informed DNN, the activations of which are deterministic functions of the DNN’s inputs. These voxelwise encoding models could predict substantial out-of-sample variance in voxel responses in all participants (block permutation test w/ 𝑅^2^-max correction: FWER-corrected *p* = 0.0002 for all subjects, which is the lowest *p*-value our permutation test could obtain). To visualize the model parameters interpretably, we divided the DNN’s inputs into three feature spaces—joint positions, joint velocities, and deviation from target positions—and computed Shapley values for each feature space, which quantify the relative contribution of each feature space to (per-voxel) predictions on the test set (Lundberg and Lee, 2017). As seen in Figure 3, the broad spatial extent of cortical activity predicted by the encoding model is relatively similar across subjects—showing recruitment of motor, somatosensory, and visual cortices—but the fine-grained encoding properties of individual voxels show little alignment across subjects despite generalizing across fMRI runs within a subject. All of this detail, though reliable within each participant, would be lost to group-averaging.

### Modeled inverse dynamics representations predict voxels in early visual cortex

While the inverse dynamics DNN-informed model outperforms chance prediction in much of cortex, this is insufficient evidence that the DNN’s representations are uniquely good predictors of voxel responses. Another explanation is that the inputs to the DNN—parametrizing the sensorimotor task state—or any nonlinear projection of those input features (of which the DNN activations are just one random choice) are sufficient to predict voxel activity. To this end, we fit a control encoding model, in which a separate kernel ridge regression (which can learn linear or nonlinear functions) is used to predict voxel activity from just the input layer to the neural network. This control model also explains substantial variance in all subjects (FWER-corrected *p* = 0.0002 for all subjects) but with no inverse dynamics prior. To isolate voxels that are uniquely well-predicted by the inverse dynamics DNN’s representations, we subtracted the variance explained (i.e., out-of-sample 𝑅^2^) by the control model from that explained by the DNN-informed model in each voxel (see Figure 3). After correcting for multiple comparisons, we find that modeled inverse dynamics representations improve prediction (paired block permutation test w/ TFCE correction: FWER-corrected *p* = 0.0044) in small patches of, strikingly, early visual cortex (V1, V2, and V3, see Fig. 4). Notably, subjects could not see their hands while their heads were in the MRI scanner, so visual monitoring of their hand cannot explain the visual cortical location.

**Figure 4:**
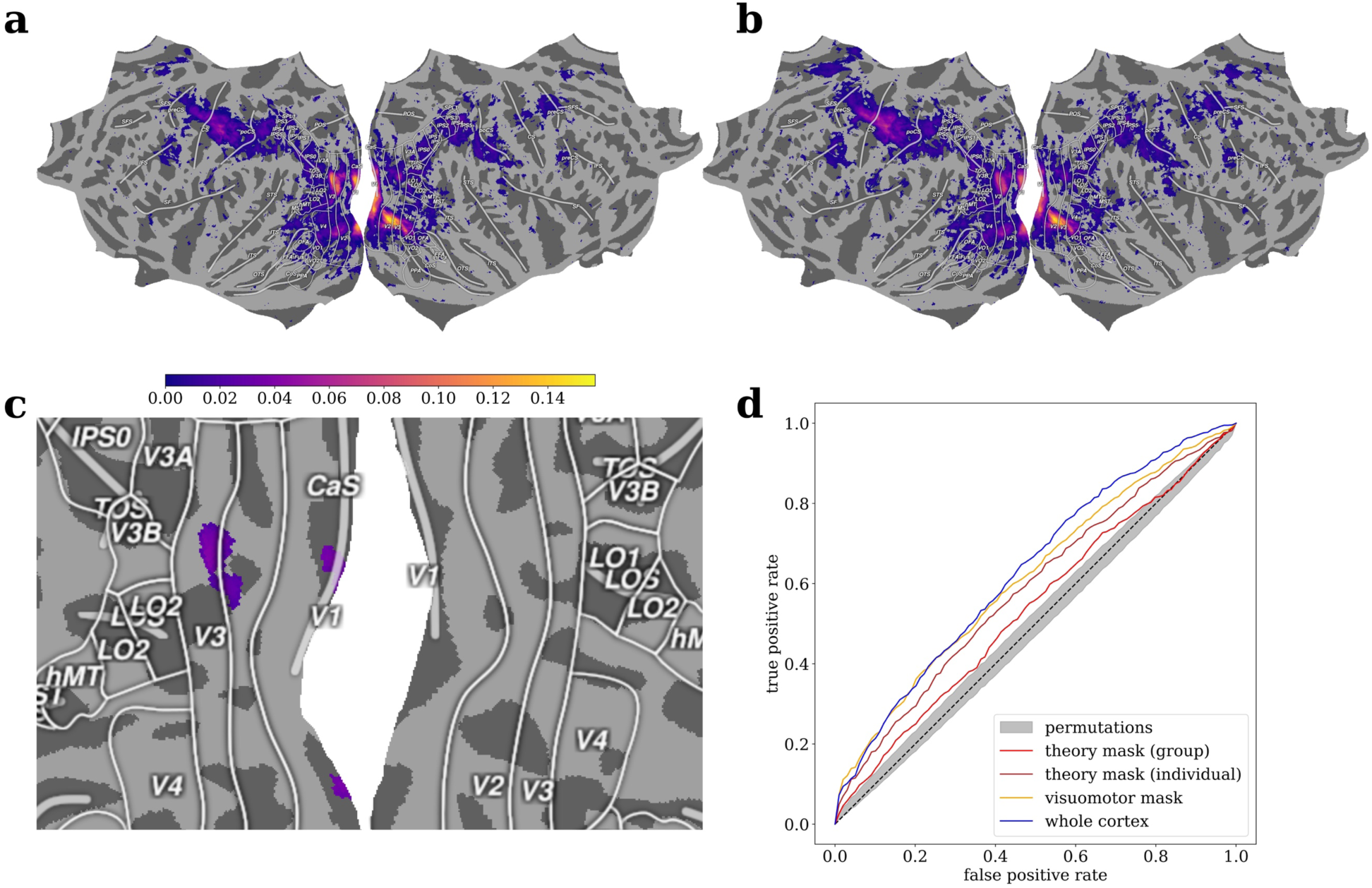
Inverse dynamics model representations predict activity in visual cortex voxels, from which sense of agency (SoA) during muscle stimulation can be decoded days later. (a) Group mean out-of-sample 𝑅^2^ for the inverse dynamics encoding model; colored voxels are where predictive performance exceeded chance after a false discovery rate correction (FDR < 0.05). (b) Group mean 𝑅^2^ for the control encoding model, also with FDR correction. (c) Voxels where inverse dynamics encoding model predicts voxelwise brain activity significantly better than chance after familywise error rate correction (FWER < 0.05). Such voxels, interpreted as involved in implementing inverse dynamics computation, were found in early visual cortex, i.e. V1, V2, and V3. (d) Receiver-operator characteristic curves, the area under which measures the sensitivity of a classifier, for decoding participants’ subjective agency judgments from inverse dynamics coding voxels (“theory mask”) using both the group-mean localization and individual-specific localizations, as well as from the full area in which encoding models could predict voxelwise activity above chance (i.e. “visuomotor”) and from the whole of cortex. The middle 90% of the permutation null distribution of the curves is shown in grey.

### Inverse dynamics responsive voxels predict sense of agency over electrically-actuated muscle movements

All participants were, similarly, preempted by FES on the overwhelming majority of trials, as indicated by the fact that “response” times—i.e., the time of the button press—are a linear function of the stimulation latency (see Figure 2b). Participants’ agency judgements were sensitive to the FES latency (mixed effects logistic regression: beta = 20.3, *p* = 8.33e-24), indicating the latency threshold at which participants reported causing ∼50% of muscle movements was relatively stable within an fMRI run (though, as evident in Fig. 2, could vary substantially over time/across runs). To decode sense of agency (SoA) over these FES-induced movements from the brain, we used logistic lasso-PCR (Wager et al., 2011) to predict single-trial agency judgments from voxel activity within (1) just the inverse dynamics selective voxels identified by our encoding models, called the “theory mask” as our hypotheses predicts strong decoding from these, (2) all voxels predicted by the control encoding model, called the “visuomotor mask” reflecting cortical areas involved in the control of the hand, and (3) all of the voxels in cortex, which serves as a rough upper bound or “noise ceiling” on the potentially decodable information available in cortex.

Decoding of agency judgments from the group-level theory mask outperformed chance sensitivity (AUROC = 0.538; permutation test: *p* = 0.0006), even when controlling for the latency of the FES that induced the movement (mixed effects logistic regression: beta = 0.067, *p* = 1.71e-5). Since, other than latency, FES was identical across all trials for a participant, this indicates that the signal decoded from the theory mask explains variance in subjective SoA beyond that which could be explained by the objective muscle stimulation characteristics. However, decoding from the visuomotor mask outperformed the theory mask substantially (AUROC = 0.614; permutation test: *p* = 0.0002), as did decoding from the whole of cortex (AUROC = 0.631; permutation test: *p* = 0.0002); whole-cortex decoding nominally, but not quite significantly, outperformed the visuomotor mask (permutation test: *p* = 0.0536). Full ROC curves for each decoder can be seen in Figure 4. Since the theory mask contains those voxels our localizer indicates are best explained by the representations of an inverse dynamics model for biomechanical control, these results indicate the neural substrates of biomechanical control and of the subjective experience of agency are indeed *overlapping*, but the identified inverse dynamics coding areas cannot fully explain SoA. At least at the group level, however, most information predicting SoA seems to be contained within the visuomotor areas active in our localizer task.

In the three participants for whom our localizer was sufficiently sensitive to identify inverse dynamics selective voxels within each individual participant (see Fig. 5 and *Participant-level Results* below), using those individual-specific localizers improved decoding performance from the theory mask from an average (among these three) AUROC of 0.586 to 0.634 (see Fig. 4d), and that improvement was statistically significant (paired permutation test: *p* = 0.0016). In fact, this increase was substantial enough that, using individual-specific localizers, the theory mask nominally outperformed decoding from the visuomotor (AUROC = 0.595) and whole-cortex (AUROC = 0.630) masks. While the participant-specific inverse-dynamics localizations varied surprisingly widely even while always residing within early visual cortex (see Fig. 5), the observed performance boost suggests such heterogeneous results may represent meaningful individual variation—consistent with recent suggestions that brain structure-function organization may be more variable across individuals than previously thought (Albantakis et al., 2024; Nakuci et al., 2024).

**Figure 5:**
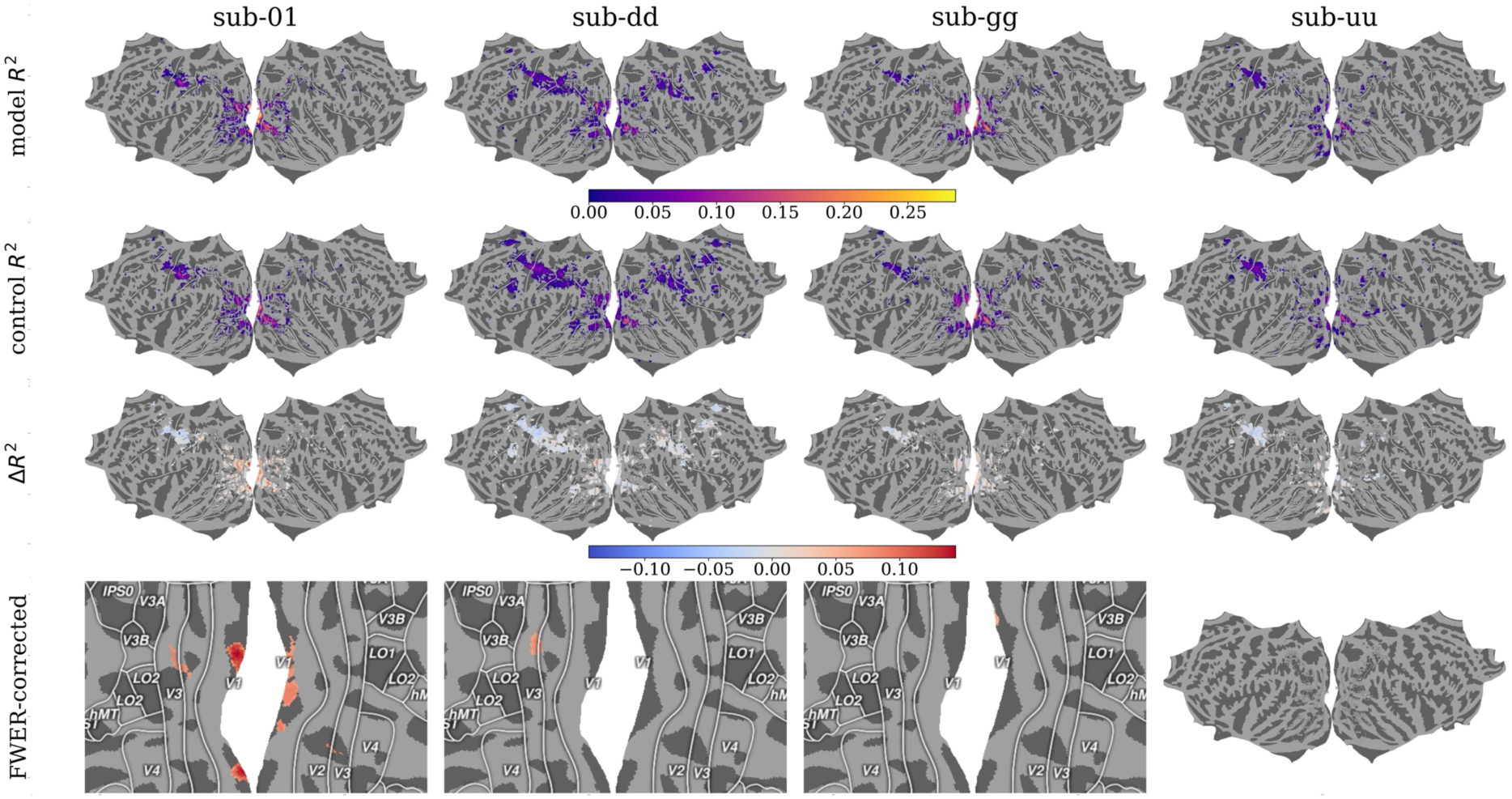
Participant-level encoding model performances. Out-of-sample variance in voxelwise activity explained by the control model was subtracted from that explained by the DNN approximating an inverse dynamics model. Voxels significantly better explained by inverse dynamics model representations, after correcting for multiple comparisons (FWER < 0.05), are interpreted as encoding similar representations.

### Participant-level Results and Prevalence Inference

While our inverse-dynamics voxelwise encoding model did predict out-of-sample BOLD data throughout a wide swath of visuomotor areas in all participants (block permutation test w/ 𝑅^2^-max correction: FWER-corrected *p* = 0.0002 for all subjects), it only outperformed the control model in three out of four subjects (paired block permutation test w/ TFCE correction: FWER-corrected 𝑝_01_= 0.0006, 𝑝_𝑑𝑑_ = 0.0300, 𝑝_gg_ = 0.0494, 𝑝_𝑢𝑢_ = 0.946; 𝑝_all_ = 0.0005). The exact location of identified cortical areas varied across those participants but was always contained to early visual cortex (V1, V2, V3; see Fig. 5). These results satisfy the sample-size sufficiency criteria of Schwarzkopf and Huang (2024), as the sequential probability ratio test (SPRT) would conclude in favor of the inverse-dynamics encoding model outperforming the control encoding model at least somewhere in the brain assuming any within-participant sensitivity—or the statistical power of the localizer within a participant—in the range [0.138, 1). A Bayes Factor estimated using *p*-curve mixture models agrees with SPRT that it is substantially more likely our localizer is sensitive to inverse dynamics selective voxels in all of, rather than none of, the population (as 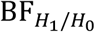 = 10.89; i.e. the alternative hypothesis explains the observed sequence of *p*-values 10.89 times better than the null hypothesis). However, the evidence leans toward an explanation where the sampled population is a mixture of people who show the tested effect and those who do not (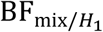 = 3.86 and 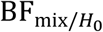 = 42.03); in other words, the data moderately support the explanation that failure to detect an individual difference in sub-*uu* is due to a real individual difference rather a lack of statistical power. Assuming the population is indeed heterogenous, we would expect the prevalence of subjects on whom the inverse-dynamics localizer would be sensitive to be about 61.0% (95% HDI: [20.2%, 97.1%]) of the sampled population.

Our main hypothesis test entails decoding SoA from voxels selectively predicted by inverse dynamics representations. Since we were unable to identify any such voxels in sub-*uu* (whether due to insufficient power or to an individual difference), this subject is not included in individual-level decoding analyses even though they were included in the group-level analysis. For the remaining participants, model weights for all decoders and their ROC curves are visualized in Figure 6. SoA could be decoded from the theory mask with above-chance *accuracy* in all three subjects (accuracy_01_ = 0.561, accuracy_𝑑𝑑_= 0.546, accuracy_gg_ = 0.599; permutation test: 𝑝_01_ = 0.0006, 𝑝_𝑑𝑑_ = 0.0266, 𝑝_gg_= 0.0002; 𝑝_all_ = 6.7e-7). The theory mask also nominally predicted SoA with above chance *sensitivity*, our preferred decoding metric, in all participants (AUROC_01_ = 0.600, AUROC_𝑑𝑑_ = 0.521, AUROC_gg_ = 0.638), but this was only significant for two subjects (permutation test: 𝑝_01_= 0.0002, 𝑝_𝑑𝑑_= 0.167, 𝑝_gg_= 0.0002; 𝑝_all_ = 1.3e-6). Here again, the SPRT would have concluded that voxels representing inverse dynamics contain decodable information about subjective agency prior to our sample size, assuming any effect size such that within-participant sensitivity is in the range [0.245, 1) (Schwarzkopf and Huang, 2024). And again, Bayes Factor evidence agrees resoundingly with the SPRT that it is more likely this effect—an overlapping cortical substrate of actual biomechanical control and of subjectively experienced agency—is present in the whole population than in none of the population (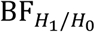 = 2,792.98). However, this time our Bayes Factor analysis indicated the data would be more parsimoniously explained by a population in which all subjects show the effect (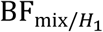 = 0.598), in which case the failure to reach significance in sub-*dd* would be considered a false negative.

**Figure 6:**
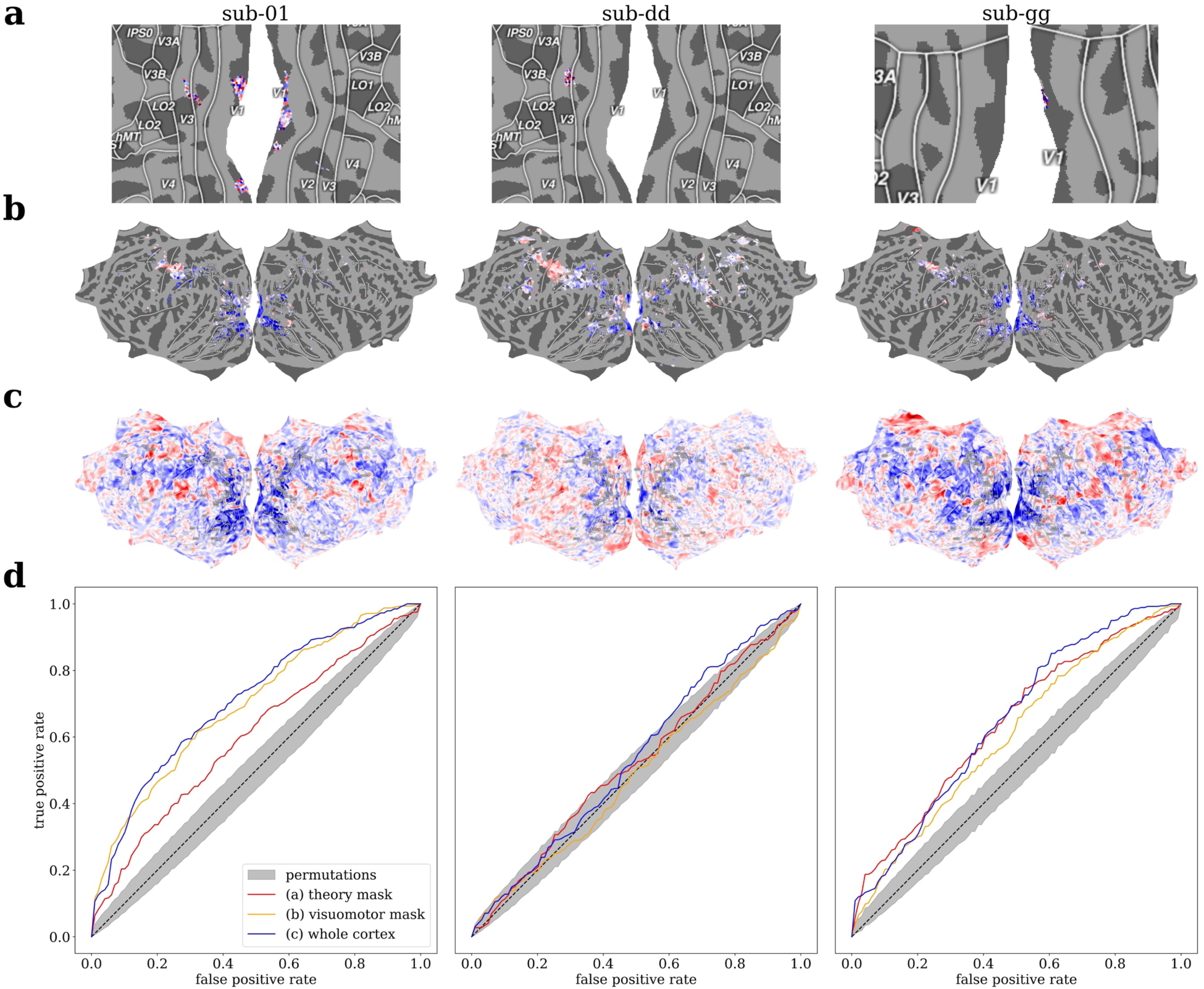
Participant-level decoding of SoA during muscle stimulation. Linear model weights for classifying between stimulation-evoked muscle movements that subjects erroneously reported were self-caused from those they did not, scaled to be between −1 and 1, are shown for (a) the “theory mask” containing only voxels that were best predicted by inverse dynamics model representations during the motor task days earlier, (b) a “visuomotor mask” containing all voxels that were predicted above-chance by the control encoding model, representing all voxels putatively involved in hand visuomotor control, and (c) all of cortex. (d) Receiver-operator characteristic curves illustrating the cross-validated performance of the three decoding models, compared to a null distribution generated by shuffling the test-set labels. The theory-driven models exceed chance sensitivity in the same two subjects whose SoA judgments were behaviorally sensitive to the latency of muscle stimulation, and they exceed chance accuracy in all three subjects.

## Discussion

The fact that our inverse dynamics encoding model successfully selected a set of voxels from which sense of agency (SoA) could be reliably decoded days later is consistent with our hypothesis that SoA is a conscious indicator that the brain can formulate a control policy for the current sensorimotor context. More conservatively, our results broadly support the idea that our internal representations for implementing motor control—not just related prediction errors (Haggard, 2017)—are involved in generating SoA.

That voxels best explained by our inverse dynamics model were found in early visual cortex (EVC) is surprising. In retrospect, there was reason to suspect early sensory areas would be involved in inverse dynamics computation. We now know encoding of movements is not restricted to canonical motor areas but ubiquitous throughout (especially sensory) cortex (Musall et al., 2019; Stringer et al., 2019). Hand movement type can be decoded from human EVC prior to movement initiation (Monaco et al., 2020), hand-selective visual regions appear to reflect potential grasp movements when viewing 3D tools (Knights et al., 2021), and reach direction can be decoded from EVC even in congenitally blind subjects (Bola et al., 2023). While early proposals suggested inverse dynamics computation would occur in motor cortex itself (Schweighofer et al., 1998), subsequent experimental evidence has long suggested such computation begins upstream of premotor cortex in visuomotor pathways (Ghasia et al., 2008; Xivry and Ethier, 2008). There is control-theoretic justification for some inverse dynamics computation to occur near primary sensory areas; in robotics, fast but “suboptimal” control policies routinely benchmark better than theoretically optimal but slower policies, as the benefits of more rapidly updating (imperfect) actions given feedback seem to outstrip that of step-by-step optimality (Howell et al., 2022). As such, there may well be a performance benefit to beginning inverse dynamics computation (i.e. movement planning) as early in sensory-to-motor pathways as possible.

It is not obvious whether the localization of inverse dynamics or sense of agency related representations to early visual cortex would generalize to other modalities. Since hand movements are decodable from EVC even in congenitally blind subjects (Bola et al., 2023), it may well be that inverse dynamics computations for the hand are present in EVC even when the relevant sensory information is arriving through another modality (e.g. auditorily). However, it is less clear how much the cortical locus of inverse dynamics computation, or sense of agency for that matter, should be expected to vary by effector; both tasks in this study involve only movements of the hand. We suspect that the same result—that sense of agency can be decoded from the same cortical areas that encode inverse dynamics—should hold regardless of whether the locus of inverse dynamics computation varies across effectors, just as our results seem to suggest that the locus meaningfully varies (within EVC) across our four participants (i.e. decoding of agency is better in Session 2 when using participant-specific inverse dynamics ROIs identified from Session 1). For now, though, this is only speculation.

Importantly, however, the fact our specific localization approach did not highlight canonical motor areas as unambiguously involved in implementing inverse dynamics should not be mistaken as suggesting they are *not* involved, which would be biologically implausible. Firstly, the representations of the DNN we use to approximate an inverse dynamics model *do* predict brain activity across a wide array of areas in the dorsal visual pathway and somatomotor cortex (see Fig. 4a). However, since these areas are also quite well—even better—predicted by the control encoding model, our computational approach cannot rule out alternative explanations for high encoding model performance. This is an expected limitation of our approach. For example, the output of our DNN inverse dynamics model is a vector of muscle activations, which should be reflected in the corticospinal output generated in motor cortex. However, only a very small fraction of neurons in primate motor cortex resemble corticospinal output; the overwhelming majority of the motor cortical activity is accounted for by the network-level dynamics necessary for generating muscle activity rather than the muscle activity itself (Saxena et al., 2022; Churchland and Shenoy, 2024), and thus the aggregate population response is dominated by action-nonspecific features such as movement timing (Kaufman et al., 2016). Our control encoding model can easily learn such features from our data, while our inverse dynamics model was constrained to an *a priori* set of features—so it is no surprise the control model performs better when predicting motor cortical fMRI, a reflection of aggregate neural activity. In contrast, a very similar approach to our own applied with single-cell resolution in a rodent model does indeed revealed inverse dynamics representations in motor cortex (Aldarondo et al., 2024). Aldarondo et al. (2024) additionally identified cells in their other recording site, the sensorimotor striatum, as encoding inverse dynamics, and there is a general consensus that some inverse dynamics computation likely occurs in the cerebellum (Wolpert et al., 1998); neither of those subcortical regions were investigated in the present study, since both regions are often argued not to directly influence conscious awareness—our main focus (Koch, 2018). In other words, our approach should be considered only a *partial* localizer, with the highlighted regions (i.e. V1, V2, V3) representing only a subset of the larger network involved in inverse dynamics computation—in particular, the subset of cortical regions in which prediction of neural activity could not be explained by potentially confounding factors that would be captured by the control model, such as encoding of the participants’ actual movements or of the presented target hand positions.

We also note the it is extremely likely the DNN we used to approximate an inverse dynamics model does not fully capture how humans solve the inverse dynamics problem. Integrating the rapidly advancing state-of-the-art in artificial biomechanical control into the analysis of real neural data, as we demonstrate as a possibility in this work, promises to ameliorate this gap. The new biomechanical control focused competition track at *NeurIPS* receives dozens of submissions a year, each a computational model of inverse (and sometimes also forward) dynamics for biomechanical control (Caggiano et al., 2023, 2024); some models have been recognized in the neuroscience literature as organically reproducing key features of human biological motor control without using any human data (Chiappa et al., 2024). A long term goal of this research program is to build biologically realistic models of the motor system from the ground up (Scott, 2024); we believe more brain-like mechanistic models will also better predict real neural data, just as more performant language models have improved prediction of human brain activity during language comprehension (Antonello et al., 2024; Goldstein et al., 2024). Unlike modern language models, however, which are explicitly trained to mimic human language using large volumes of human data, the DNN model we use here has never seen any human data; its representations were learned entirely from (simulated) sensorimotor experiences. Such models, then, can only learn more human-like behavior and more brain-like representations by changing the structure of the model (e.g. by changing computational architecture or imposing biological constraints) or by exposing it to a different set of sensorimotor experiences—both of which are informative for understanding the structure and function of the nervous system, furthering the long-term goal of creating mechanistic models that explain observed biological data. However, in the near term, we recognize constraining the feature space of our encoding model to that learned by a neural network independent of any human data limits predictive performance.

Modeling cortical control of the musculature with such granularity has only recently become feasible. While motor control is a well-developed field with numerous empirically successful computational models, such models are usually specified at the level of macroscopic movements or net force output instead of that of musculoskeletal kinematics and kinetics—and for good reason. Embodied control is too high-dimensional a problem to specify model parameters by hand, and biomechanical simulation has been too slow to learn parameters with reinforcement learning. Just recently, it became possible to port existing biomechanical models into robotic-grade physics simulators (Wang et al., 2022). Now, a growing library of simulation environments features not just healthy musculoskeletal systems but clinically informative cases incorporating sarcopenia, tendon transfer surgeries, and assistive exoskeletons (Caggiano et al., 2022). To capitalize on such computational advances, it is critical to model neural activity at the level of individual subjects rather than directing all resources to group studies (Naselaris et al., 2021); to this end, the present work demonstrates that robust brain-model relationships can be detected not just at the group level but also at the level of the individual. Even healthy brains show variation in functional organization (as in Fig. 3), but neuroscience promises to improve the lives of those whose neural responses may look *least* like the average brain—with neurological, motor, and musculoskeletal pathologies. The present work demonstrates that biomechanically detailed control models can be used to predict human brain activity, affording investigation of specific relationships between motor system computations, behavior, and subjective experience.

## Author Contributions

**A.F.C.:** Software and Writing - review & editing. **H.C.N.:** Conceptualization, Funding acquisition, Resources, Supervision, and Writing - review & editing. **J.P.V.:** Conceptualization, Data curation, Formal analysis, Funding acquisition, Investigation, Methodology, Project administration, Software, Validation, Visualization, Writing - original draft, and Writing - review & editing. **P.L.:** Conceptualization, Funding acquisition, Resources, and Writing - review & editing. **R.N.:** Resources and Writing - review & editing.

## Acknowledgements

This work was supported by NSF BCS 2024923 to H.C.N. and P.L., and J.P.V. was supported by NSF GRFP DGE 1746045. Data acquisition was completed with equipment funded by NIH S10OD018448 to the Magnetic Resonance Imaging Research Center at the University of Chicago, and analysis used resources provided by the University of Chicago’s Research Computing Center.

